# Predicting synaptic connectivity for large-scale microcircuit simulations using *Snudda*

**DOI:** 10.1101/2021.04.15.439985

**Authors:** J J Johannes Hjorth, Jeanette Hellgren Kotaleski, Alexander Kozlov

## Abstract

Simulation of large-scale networks of neurons is an important approach to understanding and interpreting experimental data from healthy and diseased brains. Owing to the rapid development of simulation software and the accumulation of quantitative data of different neuronal types, it is possible to predict both computational and dynamical properties of local microcircuits in a ‘bottom-up’ manner. Simulated data from these models can be compared with experiments and ‘top-down’ modelling approaches, successively bridging the scales. Here we describe an open source pipeline, using the software Snudda, for predicting microcircuit connectivity and for setting up simulations using the NEURON simulation environment in a reproducible way. We also illustrate how to further ‘curate’ data on single neuron morphologies acquired from public databases. This model building pipeline was used to set up a first version of a full-scale cellular level model of mouse dorsal striatum. Model components from that work are here used to illustrate the different steps that are needed when modelling subcortical nuclei, such as the basal ganglia.

## Introduction

Neuroscientists are producing data at an ever growing rate, and sharing the data in public databases. Within the computational neuroscience field, hypothesis-driven modelling has over many decades generated new ideas that in turn have been tested via experiments. Recently a data-driven mechanistic modelling approach has also gained ground thanks to new technologies allowing the collection of large quantities of useful data. In particular, large-scale spiking neural network models have been reconstructed in a data-driven manner and simulated (Markram et al., 2015; Gratiy et al., 2018; Migliore et al., 2018; Casali et al., 2019; Einvoll et al., 2019; Billeh et al., 2020; Hjorth et al., 2020). Collecting data from the brain at multiple biological scales from mouse, non-human primates, and human, are important goals of several of the big brain initiatives (Insel et al., 2013; Amunts et al., 2019; Okano et al., 2015; Grillner et al., 2016), and will further facilitate and speed up this modelling process. In parallel, various brain simulation tools have been optimized to capitalize on supercomputers (Hepburn et al., 2012; Plesser et al., 2015; Carnevale and Hines, 2006; Hines et al., 2009, Kumbhar et al., 2019; Ray and Bhalla, 2008; Gleeson et al., 2010; Jordan et al., 2020; Akar et al., 2019). In this respect, the principles of FAIR – Findable, Accessible, Interoperable, Reusable (Wilkinson et al., 2016) – are important for catalysing this process, both with regard to the experimental data, the data-driven models as well as the software used during the modelling and simulation process. We also believe that to be able to reproduce the actual model reconstruction process, given the same or new additional data, is one important aspect of the FAIR criteria when making the modelling process transparent, repeatable, reusable and comparable.

Here we present our open source modelling pipeline that facilitates a reproducible, data-driven reconstruction of cellular level network/microcircuit models. This pipeline inspired by the cortical column microcircuit (Markram et al., 2015) has been applied to predict a full-scale microcircuit model of the mouse dorsal striatum (Hjorth et al., 2020). *Snudda* is a software to create a detailed network of connected neurons, where the connectivity is derived from reconstructed neuronal morphologies as well as from more qualitative experimental knowledge (see also Reimann et al., 2015). ‘Snudda’ means ‘touch’ in Swedish, and it supports the creation of a network with connectivity based on touch detection. If detailed morphological data exist, the algorithm looks for close appositions between axons and dendrites, which are locations for putative synapses. Thus the morphology restricts where connections can be positioned. Snudda can also define the axon using a probability cloud if a reconstructed axon is missing. This is an extension of the method where the connection probability is proportional to the overlap of two spheres representing axons and dendrites (Humphries et al., 2009). Based on a set of rules, as described below, the putative synapses are then pruned to match the connectivity seen from pairwise experimental recordings, or other types of connectivity experiments. The same technique can be applied to also place gap junctions. The generated network can then be simulated using parallel NEURON (Carnevale and Hines 2006). Similar approaches have been used to build the somato-sensory cortex microcircuit (Markram et al., 2015; Colangelo et al., 2019), visual cortex model (Billeh et al., 2020; Dai et al. 2020), cerebellar network (Wichert et al., 2012; Sudhakar et al., 2017; Casali et al., 2019) and hippocampal neurons (Migliore et al., 2018).

The reconstruction of a local microcircuit model (such as striatum) consists of the following steps: a) experimental data acquisition of the electrophysiological and morphological properties of neuronal types, and also characterisation of synapses, b) optimization of neuron and synapse models, c) placement of the model neurons in the brain volume to be modelled, d) prediction of microcircuit connectivity *in silico*, e) constraining and emulating inputs for the model, and finally f) simulating the microcircuitry. Our software Snudda is used for steps c)-f). Software Treem for improving the morphological reconstructions in preparatory step b) is described at the end. The code is publicly available on GitHub (https://github.com/Hjorthmedh/Snudda/) and (https://github.com/a1eko/treem). Below we will go through the different steps and provide code to set up an example network using Snudda, followed by explanation of the configuration files, network building and simulation process, as well as some preprocessing options. The network example corresponds to a 0.5 mm cube within the mouse striatum. We will assume that we already have a set of electrophysiologically optimized neurons and synapses, e.g. using the optimization tool BluePyOpt (Van Geit et al., 2016). Examples of neuron and synapse models relevant for striatum are provided on GitHub (in the `snudda/examples` folder with scripts and notebooks).

Our approach offers novel contributions in the following aspects: (i) we design and present a complete, free and open source toolchain for building and simulating anatomically constrained biologically detailed neural networks including morphology-based neuron touch detection; (ii) we illustrate the use of this platform on the example of the striatal microcircuit, implemented at a very detailed level and accuracy; (iii) we include all tools and parameters in the source code repository, enabling other labs to reproduce as well as reconstruct our striatal model with new data when it becomes available.

### Getting started with Snudda

Snudda is available for download from GitHub (https://github.com/hjorthmedh/Snudda) or from PyPi through pip3 install snudda. Both source code and the data files necessary to set up a striatal network are provided, Table 1 provides an overview of the *Snudda* directory structure. Snudda is compatible with Linux, Mac and Windows 10.

**Table 1.** Snudda directory structure.

|  |  |
| --- | --- |
| snudda | Snudda directory, code usually executed from this folder |
| snudda/data | Snudda data folder |
| snudda/data/mesh | 3D meshes for brain structures |
| snudda/data/neurons/striatum | morphology and parameters for striatal neurons in subfolders |
| snudda/data/neurons/mechanisms | NEURON mechanisms folder with mod-files |
| snudda/data/synapses | synapse model parameters |
| snudda/data/input_config | input configuration files |
| snudda/input_tuning | scripts for synaptic input tuning |
| snudda/plotting | scripts for plotting networks, includes subfolder with blender scripts |
| snudda/utils | scripts for small tasks |
| examples | examples on how to run snudda, contains shell scripts and notebooks |
| tests | contains unit and regression tests |
| tests/networks | networks used or created for testing |
| tests/validation | morphologies and mechanisms used for testing |

In the directory `snudda/data/neurons/<region>` there are separate subdirectories for each neuron type (`<region>` in our use case is `striatum`). Each of those directories contains multiple subdirectories, one for each unique morphology from that neuron type. The neuron directories include the morphology in SWC format, a JSON parameter file with one or more sets of optimised neuron parameters from BluePyOpt, a JSON mechanism file specifying which mechanisms are present in each compartment, and a JSON modulation file which specifies the neuron modulation of the neuron. The JSON file format was chosen as it is a standardised and human readable way to store structured data. The `neurons` folder also has a `mechanisms` folder containing the NEURON model description language `.mod` files with definitions of ionic mechanisms.

To keep the networks separate, each generated network has its own directory which contains a `network.json` file that links together all the different components that make up the network. The `network.json` file can be manually created, or in the case of the striatal network there is a way to automatically generate a `network.json` file of user specified size. The script `init.py` can be extended to create networks of other brain structures. A Jupyter notebook in `examples/notebooks` shows an alternative example for how to define brain slices and other structures.

In the main `Snudda` directory there is an `examples` folder with useful scripts and Jupyter notebooks for generating and running networks. The directory `snudda/plotting` contains scripts to plot simulation results as well as visualise the network or parts of it using Blender (https://www.blender.org/).

### Use case: striatal microcircuit

First we create an example striatal network, then further down we go through all configuration details. The `network-config.json` can be generated using the snudda init shell command. In the below example a homogeneous cube with 0.5 mm side length in the mouse striatum is generated (**Figure 1A**). This corresponds to 10,062 neurons (**Figure 1B**) using the estimated average density of striatal neurons (Rosen and Williams, 2001), but the number of neurons can be varied depending on computational resources and research questions. The neurons are taken from the `data/neurons/striatum` directory, where every neuron type has its own directory, e.g. `dspn` or `ispn`. A neuron type is represented by one or more single-cell models, each in its own subdirectory as described above. When the network is initialised, the `init.py` code will look in the folders of the different neuron types and instantiate single-cell models in random order. No modifications of the neuron models other than rotations are applied at runtime. In order to improve the cell diversity, one should populate the neuron types directories with a sufficient number of different single-cell models.

**FIGURE 1.**
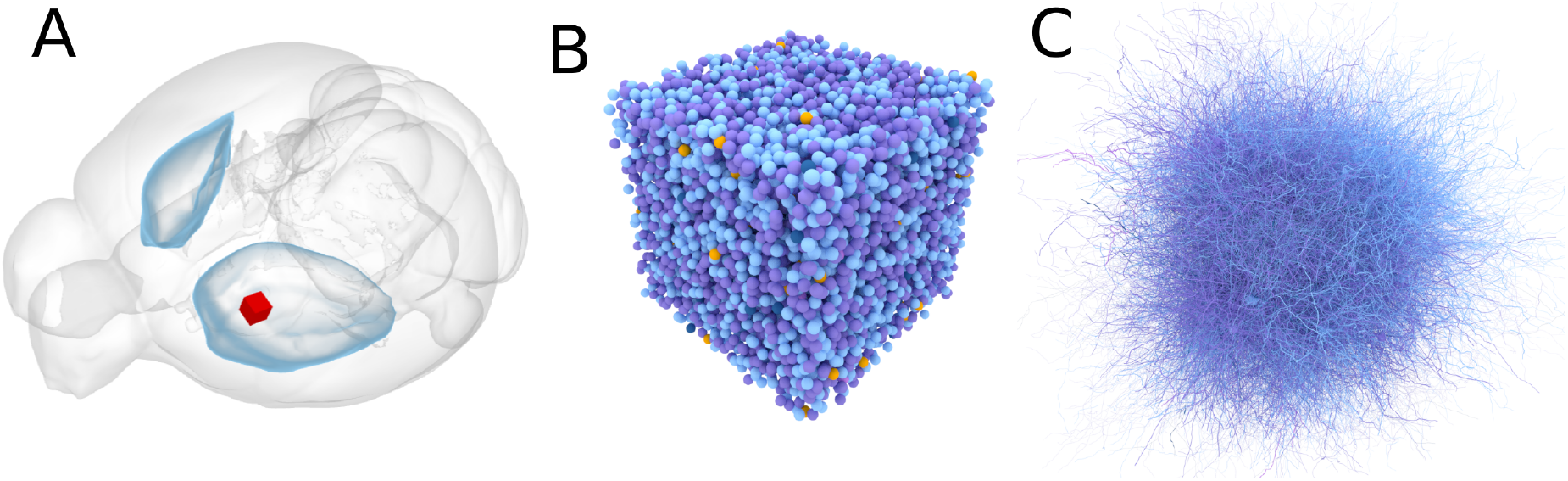
Example of the volume definition. **(A)** Selection of the volume of interest (red cube, size of the side 500 micrometers) inside the left part of the dorsal striatum (blue shells beneath the cerebral cortex). **(B)** The 10,062 neuron somas placed within the red cube. **(C)** Illustration of 100 neurons showing the complexity of axons and dendrites.

The following commands can be used in a terminal window to generate the example network that we have used for the figures in this article. We go into more detail in the sections later in the article. There are additional Jupyter notebook examples in `examples/notebooks` and `examples/Neuroinformatics2021`. The first step is to create a configuration file `network.json` specifying e.g. 10,062 neurons in a directory called `smallSim`.

~~~
simName=smallSim
snudda init $simName --size 10062 --overwrite
~~~

Next we need to place the neurons in a specified volume. For large simulations the neurons are placed inside a volume representing the mouse striatum (**Figure 1A**), while smaller networks use a simple cube (**Figures 1B,C**). This is done to preserve physiological neuron densities in the simulations. The mesh definition of the striatal volume, or other structures, can be extracted from databases such as the Allen Brain Atlas of the mouse brain. The command to place the neurons inside the volume defined by the mesh is:

~~~
snudda place $simName
~~~

The next step is the touch detection (**Figure 2A**). Here the algorithm voxelizes the space, and looks for overlaps (within a certain predefined distance) between axons and dendrites from different neurons (**Figure 3A,B**) (Hellwig, 2000).

**FIGURE 2.**
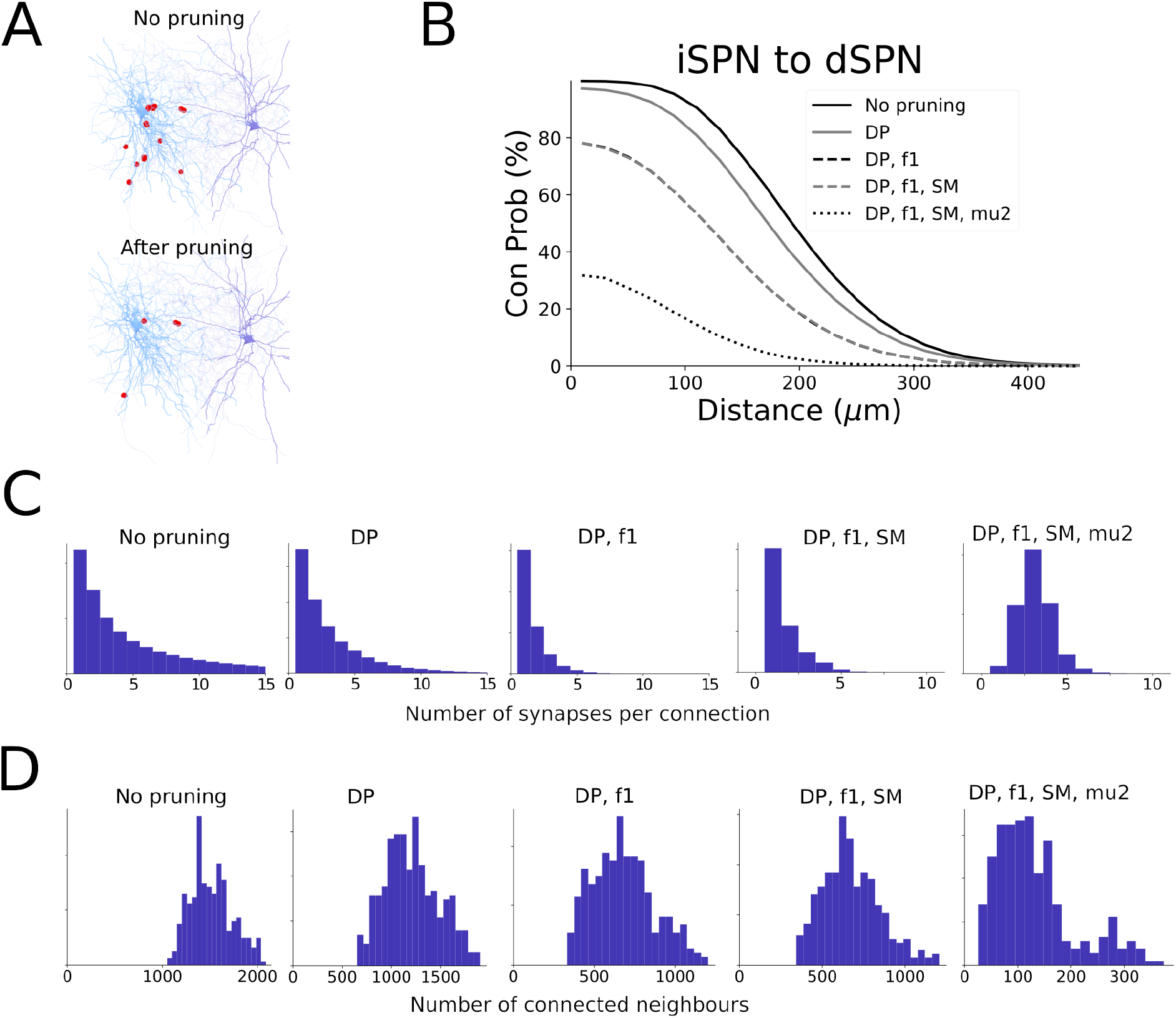
Example of the synaptic pruning procedure when connecting neurons within microcircuit. **(A)** Putative synapses detected between iSPN and dSPN shown on top, and remaining synapses after pruning shown below. **(B)** Connection probability as a function of distance after each of the pruning steps. Distance dependent pruning filters synapses based on the distance to the soma on the postsynaptic neuron. A fraction *f1* of all synapses is removed. The soft max synapse filter does not disconnect any connected pairs as it only reduces the number of synapses of pairs that are connected by a large number of synapses. Finally *mu2* filters neuron pairs with few synapses, and leads to a large reduction in connectivity. **(C)** Number of synapses between connected pairs. **(D)** Number of connected neighbours each post synaptic neuron has.

**FIGURE 3.**
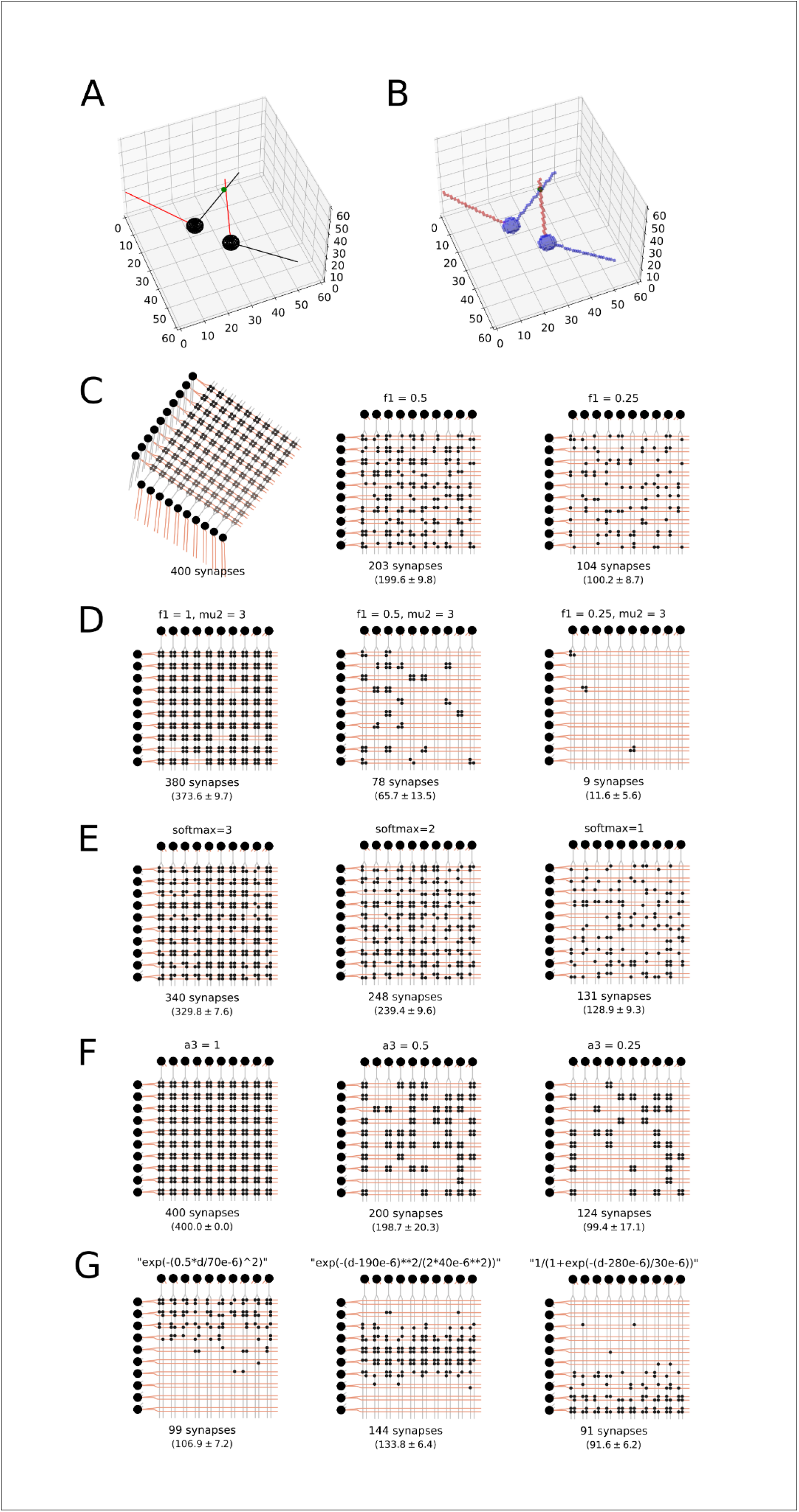
Touch detection and pruning. **(A)** Illustration of ball and stick neurons with soma and dendrites marked in black, axon in red and synapse in green. **(B)** Corresponding two neurons in the hypervoxel representation. The neurites are traced by taking small steps along *Δx, Δy, Δz* corresponding to the direction of the neurite. Where axon and dendrites occupy the same voxel a putative synapse is detected, here marked by a green dot. The volume of the soma is also voxelized. **(C)** Ball-and-stick neurons arranged so their synapses are on a grid, with four synapses connecting every neuron pair. Pruning with parameter *f1*=1, 0.5 and 0.25 keeping 100%, 50% and 25% of all synapses, respectively. Number of synapses retained shown under each network (including a numeric estimate of mean and standard deviation, *n*=1000). **(D)** Combination of *f1* and *mu2* pruning. Here *P* = *f1* · 1.0 / (1.0 + exp(-8.0 / *mu2* · (*nSynapses* - *mu2*))), where *P* is the probability to keep a synapse, *nSynapses* is the total number of synapses connecting the neuron pre-post pair. **(E)** *SoftMax* pruning step. If there are more than *softMax* synapses then the probability of keeping synapses between that pair is *P* = 2 · *softMax* / ((1 + exp(-(*nSynapses* - *softMax*) / 5)) · *nSynapses*). **(F)** Pruning *a3* = 1, 0.5 and 0.25 removing all synapses between a connected pair in 0%, 50% and 75% of the cases, respectively. **(G)** Distance dependent pruning of proximal, medial and distal synapses. Jupyter Notebooks to generate this figure are available on Snudda GitHub in the `examples/Neuroinformatics2021` folder).

~~~
snudda detect $simName --volumeID Striatum
~~~

The touch detection will create a putative set of synapses at all the close appositions between axons and dendrites. However, not all close appositions correspond to real synapses, as explained in detail further down. The next step prunes the set of putative synapses to match the connectivity seen in experimental pairwise recordings (**Figure 2B**, distance dependent connectivity). The rules used for pruning are qualitatively similar to what Markram et al. (2015) created for their cortical network. The parameters for the pruning (**Figure 3C-G**) are specified in the `network.json` file, explained more in detail below. The command to perform the pruning is:

~~~
snudda prune $simName
~~~

Code to generate figures analysing the connectivity (**Figure 2C,D**) (distance dependent connection probability, histogram showing the number of synapses between connected neighbours, histogram showing the number of connected neighbours) is in snudda/analyse_striatum.py, also see examples/Neuroinformatics2021 for Jupyter notebooks.

Next we need to generate external synaptic input for the network simulation. Here we specify how much time we want to generate inputs for. The parameters for the synaptic input from cortex and thalamus are defined in a separate JSON file:

~~~
cp -a data/input_config/input-tinytest-v9-freq-vectors.json
$simName/input.json
snudda input $simName --input $simName/input.json --time 3.5
~~~

The last step involves compiling the `.mod` files, and then running the simulation.

~~~
nrnivmodl data/neurons/mechanisms
snudda simulate $simName --time 3.5
~~~

There are two functions in the plotting directory that allow the user to plot either the spike raster or the voltage traces, `plotting`/`plot_spike_raster.py` (**Figure 4**) and `plotting/plot_traces.py`, the latter requires the user to have run the simulation with the `--voltOut` parameter to also save the voltage traces. The simulation output files are stored in the `$simName/simulations` directory.

**FIGURE 4.**
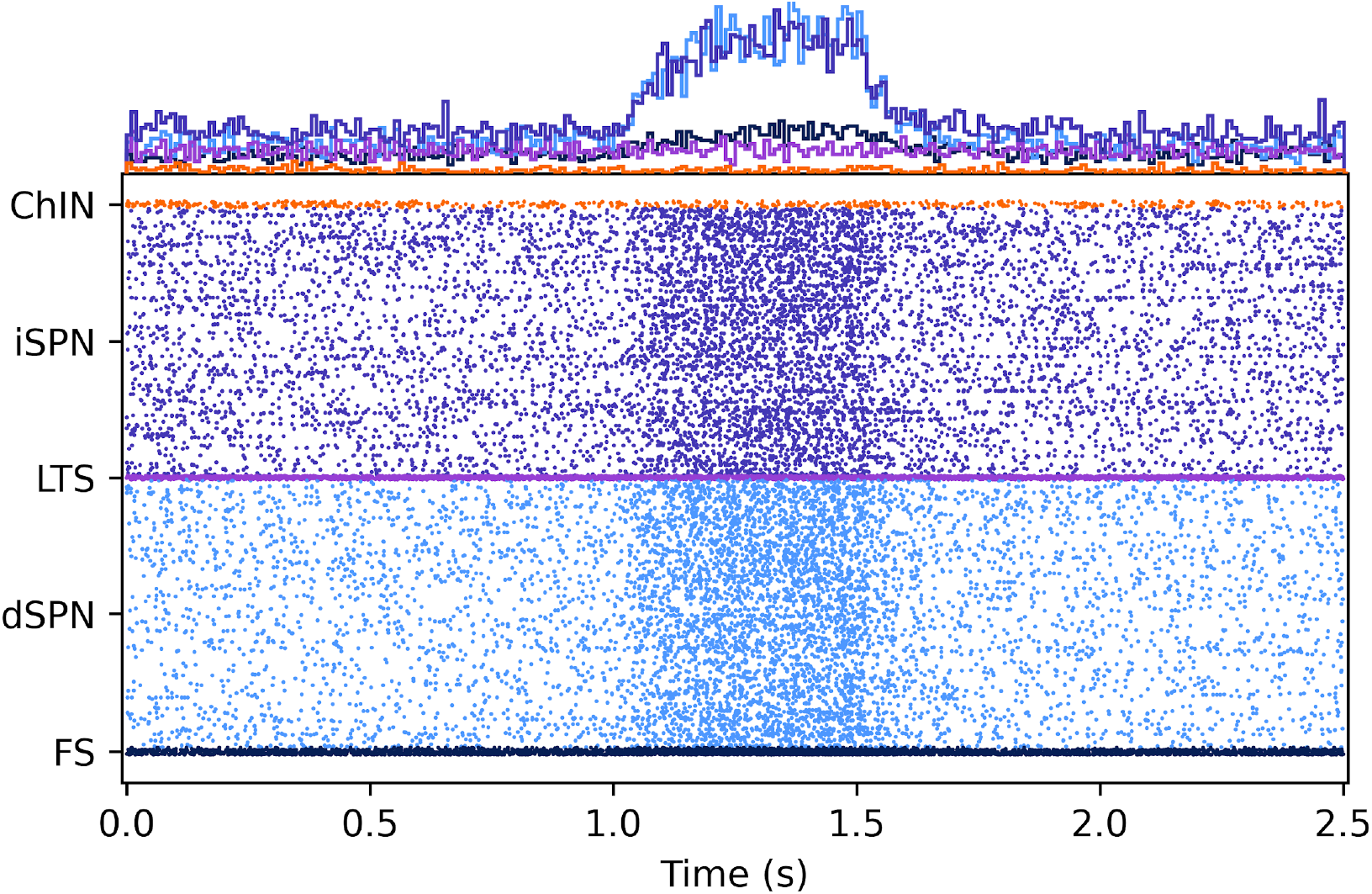
Simulation of 10,062 striatal neurons (4872 dSPN, 4872 iSPN, 133 FS, 113 ChIN, 72 LTS) receiving cortical and thalamic input. The cortical drive is increased for half a second at 1.0 seconds. The histograms above the spike raster show the total number of spikes in the respective neuron populations. (See the `striatum_example*` notebooks in the `example/notebooks` folder, note that the size of the network was changed.)

### Validation against anatomical data

The network building results in Figure 2 have been validated in our previous study. Figure 8, and S4-S7 in Hjorth et al. (2020) showed experimental pair-wise connection probability between neuron types and how the Snudda generated network matched the data, and matched estimations of number of synapses between connected pairs.

To validate the mean synaptic density in the simulated network, we use the data from the recent studies with genetic labeling and electron microscopy techniques. Santuy et al. (2020) estimate a density of 1.41 synapses/μm^3^ in the striatum, with 4.4% of the symmetric synapses (range 1.29 - 20.23%), which corresponds to 62 million GABAergic synapses per mm^3^. In our striatal model 500,000 neurons (6.2mm^3^) has 469 million intrastriatal GABAergic synapses which corresponds to 75.6 million synapses per mm^3^, well within the experimental range (18 - 285 million). This also agrees with another independent study by Cizeron et al. (2020).

### Snudda configuration explained

#### Connectivity configuration

Running the command `snudda init mysimulation --size N` generates the Snudda network configuration file `network-config.json` where `N` is the size of the network to create. The `network-config.json` contains the blueprints for the striatal network hierarchically organised into the blocks *RandomSeed, Volume, Units, Connectivity* and *Neurons*. All parameters in the configuration file are specified in SI units. Below we go into more detail how Snudda is configured.

The *RandomSeed* block specifies the random seeds used for the different steps of the network creation.

~~~
“RandomSeed”: {
  “init”: 605423731,
  “place”: 713568010,
  “detect”: 3462736139,
  “prune”: 2408118524,
  “input”: 1253064917,
  “simulate”: 3292847414
},
~~~

The *Volume* block can contain several named volumes, such as for example *Striatum, Cortex, Thalamus*. For each volume block we define *type* (e.g. *mesh*), *dMin* (minimum distance between somas in the volume), *meshFile* (this specifies the Wavefront OBJ file that defines the mesh enclosing the volume) and *meshBinWidth* (voxelization size of the mesh for determining what is inside and outside the mesh during cell placement). Future versions of Snudda will allow for density variations within the volume and directional gradients for the neurons.

~~~
“Volume”: {
  “Striatum”: {
    “type”: “mesh”,
    “dMin”: 1.5e-05,
    “meshFile”: “striatum.obj”,
    “meshBinWidth”: 5e-05
  }
},
~~~

The next block is the optional *PopulationUnits* block, which allows us to define network heterogeneities within the volumes These units may receive common input which is different from the input to other neurons. Snudda also supports different connectivity within and between the population units. These could mimic functional units that have been organised due to developmental processes or learning, e.g. a functional unit within the striatum could map to an ‘action channel’ (Gurney et al., 2001; Lindahl and Hellgren Kotaleski, 2016; Berthet et al., 2016). There are currently two methods supported; *random* which randomly picks a fraction of the neurons for each population unit, or *radialDensity* where a density function placed in the volume determines the neurons in each population unit, see the examples on GitHub (https://github.com/Hjorthmedh/Snudda/examples/notebooks/).

An example block for *PopulationUnits* looks as follows. Here two units are defined: *UnitID* 1 with 20% of *dSPN* and *iSPN* neurons in *Striatum*, and *UnitID* 2 with 30%.

~~~
“PopulationUnits”: {
   “AllUnitID”: [1, 2],
   “Striatum”: {
      “method”: “random”,
      “fractionOfNeurons”: [0.2, 0.3],
      “unitID”: [1, 2],
      “neuronTypes”: [
         [“dSPN”, “iSPN”],
         [“dSPN”, “iSPN”]
      ],
      “structure”: “Striatum”
   }
}
~~~

In the *Connectivity* block we define the rules guiding how the different neuron populations are connected together. Each connection pair has its own block (e.g. “iSPN,dSPN”). In the example below this is illustrated with the iSPN-dSPN pair (indirect-pathway and direct-pathway striatal projection neurons, respectively), which are connected by GABA synapses.

~~~
“Connectivity”: {
   “iSPN,dSPN”: {
      “GABA”: {
         “conductance”: [
            2.4e-10,
            1e-10
         ],
         “channelParameters”: {
            “tau1”: [
               0.0013,
               1000.0
            ],
            “tau2”: [
               0.0124,
               1000.0
            ],
            “failRate”: 0.4,
            “parameterFile”:
“$DATA/synapses/striatum/PlanertFitting-ID-tmgaba-fit.json”,
            “modFile”: “tmGabaA”
      },
      “pruning”: {
            “f1”: 0.3,
            “softMax”: 4,
            “mu2”: 2.4,
            “a3”: 1.0,
            “distPruning”: “1-exp(-(0.4*d/60e-6)**2)”
      },
      “pruningOther”: {
            “f1”: 0.3,
            “softMax”: 4,
            “mu2”: 2.4,
            “a3”: 1.0,
            “distPruning”: “1-exp(-(0.4*d/60e-6)**2)”
      }
   }
},
~~~

Each connection has a *conductance* parameter, which specifies the mean and standard deviation of the conductance. The *channelParameters* gives flexibility by specifying a dictionary with the channel specific parameters, in this case it is *tau1, tau2, failRate* (the synapse failure rate) that are passed directly to the NEURON channel model. Next *parameterFile* (JSON file with additional channel parameters) and *modFile* (NEURON channel `.mod` file). The final two blocks *pruning* and *pruningOther* specify the pruning parameters for neurons within the same population unit, and neurons in different population units. The pruning parameters help shape the connectivity by parameterising the rules that define which putative connections should be removed, and which ones should be kept. The probability to keep a synapse is equal to the product of the individual pruning steps: 

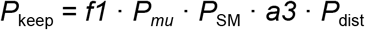

The examples given in **Figure 3C-G** are synthetic, their purpose is to illustrate pruning rules. The *f1* parameter defines how large a fraction of the putative synapses should be kept, a value of 1.0 or None means that this pruning step is not used (**Figure 3C**). For *f1*=0.5 we would expect on average 0.5 · 400 = 200 synapses kept, and for *f1*=0.25 we expect 0.25 · 400 = 100 synapses (*c*.*f*. 203 and 104 synapses randomly selected for *f1*=0.5 and *f1*=0.25, respectively, in **Figure 3C**).

The *mu2* defines a sigmoid curve used to decide whether to keep or remove all synapses between a coupled pair of neurons (**Figure 3D**): 

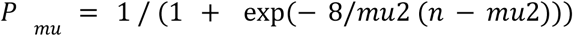

With *mu2* = 3, we have *P*_*mu*_(*n*=4) = 93.5%, *P*_*mu*_(*n*=3) = 50%, *P*_*mu*_(*n*=2) = 6.5%, *P*_*mu*_(*n*=1) = 0.5%. Thus for *f1* = 1 we expect 400 · 0.935 = 374 synapses left. For *f1* = 0.5 we have two parts to the pruning, first *f1* where half the synapses are removed, then *mu2* which operates on all the synapses between a coupled pair. We thus look at the 100 possible neuron pairs. The expected number of synapses 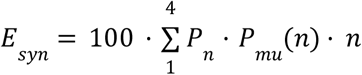, with *P*_*n*_ the probability of a neuron pair having *n* synapses after the *f1* pruning. We then get 

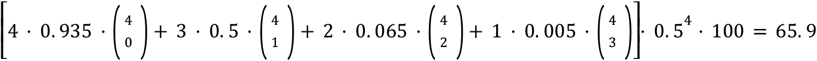

synapses, and for *f1* = 0.25 we expect on average 

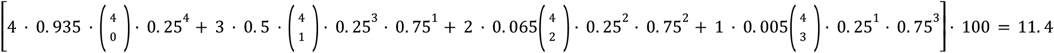

synapses.

The *softMax* specifies at which value we start applying a soft cap to the total number of synapses between the pair: 

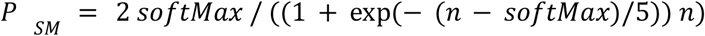

where *P*_SM_ is probability to keep a synapse, and *n* is the initial number of synapses between the pair of neurons (**Figure 3E**). For *n*=4 and *softMax*=3 yields *P*_SM_= 82.5% resulting in on average 400 · 0.825 = 330 synapses left. For *softMax*=2 around 239 synapses will remain, and for *softMax*=1 on average 129 synapses are kept.

The *a3* parameter specifies which fraction of all connected pairs to keep, e.g. 0.8 means that 20% of all connected pairs will have all their synapses removed (**Figure 3F**). Here *a3* = 1, 0.5 and 0.25 result in on average 400, 200 and 100 synapses left, correspondingly. The *distPruning* defines a distance *d* dependent function *P*_dist_ : *d* ⟶ [0,1], where *d* is the distance from the soma along the dendrites (**Figure 3G**). The expected number of synapses for the distance dependent pruning is given by 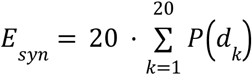 where *P*(*d*) is one of the equations in Figure 3G. In this example the distances to the soma are *d* = 56, 65, 86, 95, 116, 125, 146, 155, 176, 185, 206, 215, 236, 245, 266, 275, 296, 305, 326 and 335 μm (with 20 putative synapses at each distance). The expected number of synapses in the three cases are thus 107, 134 and 91, respectively.

Continuing our look at the network configuration file structure, the final block *Neurons* defines the different neuron populations. Here each neuron template has its own block. For each neuron template we specify four files: *morphology* (a SWC file defining the soma, axon and dendrites), *parameters* (neuron parameters optimised using BluePyOpt), *mechanisms* (NEURON mechanisms), *modulation* (a JSON file defining the neuromodulation). The template can be used to define multiple neurons, the number defined by *num*. The *hoc* parameter is optional, and intended to be used in the future when exporting to SONATA format for use with Neurodamus (Williams et al., 2018). The *neuronType* can be either *neuron* or *virtualNeuron*, the latter can be used to define axons from other structures providing input to the striatum. The *rotationMode* lets us specify if the neurons should be left unrotated, or rotated in some manner. The *volumeID* defines which volume the neurons belong to.

~~~
“Neurons”: {
   “dSPN_0”: {
      “morphology”:
“$DATA/neurons/striatum/dspn/str-dspn-1/WT-0728MSN01-cor-rep-ax.swc”,
      “parameters”:
“$DATA/neurons/striatum/dspn/str-dspn-1/parameters.json”,
      “mechanisms”:
“$DATA/neurons/striatum/dspn/str-dspn-1/mechanisms.json”,
      “modulation”:
“$DATA/neurons/striatum/dspn/str-dspn-1/modulation.json”,
      “num”: 1218,
      “hoc”: null,
      “neuronType”: “neuron”,
      “rotationMode”: “random”,
      “volumeID”: “Striatum”
   },
~~~

The `$DATA` keyword is a shorthand for the `snudda/data` folder.

### Configuring external synaptic input

The input spikes to the network are generated as prescribed in the input configuration file `input.json` in JSON format. Below we will give a simple example of how to set up input, and there are more examples available on Github in the examples/notebooks directory.

In this example the *dSPN* will each receive 200 inputs, with 1Hz Poisson random spikes. The configuration also specifies the conductance and the *tmGlut* mod file that is used by NEURON to simulate the input synapses.

~~~
{
   “dSPN”: {
      “Ctx” : {
         “generator” : “poisson”,
         “frequency” : 1,
         “conductance” : 0.5e-9,
         “nInputs” : 200,
         “modFile”: “tmGlut”
      }
   }
}
~~~

To complement the cortical (*Ctx*) input with thalamic, add a second input block with parameters inside the *dSPN* target block and give it a name e.g. *Thalamic*. The entire dSPN population will then receive both cortical and thalamic inputs.

Only one target block is applied to each neuron. When Snudda generates input for the network it iterates through all the different neurons in the simulation and picks the most specific target block that matches that neuron. In the example below a dSPN with neuron ID 5 and morphology *dSPN_0* will match all three blocks, but the neuron ID is most specific so the “5” block will be used. A *dSPN_1* morphology neuron will only match the *dSPN* block and will use that.

~~~
{
   “dSPN”: {
      “Ctx”: {
         “generator”: “poisson”,
         “start”: [2, 5],
         “end”: [3, 7],
         “frequency”: [4, 2],
         “conductance”: 0.5e-9,
         “nInputs”: 200,
         “modFile”: “tmGlut”
      }
   },
   “dSPN_0”: {
      “Ctx”: {
         “generator”: “poisson”,
         “start”: 0,
         “end”: 10,
         “frequency”: 1,
         “conductance”: 0.5e-9,
         “nInputs”: 200,
         “modFile”: “tmGlut”
      }
   },
   “5”: {
      “Ctx”: {
         “generator”: “poisson”,
         “start”: 2,
         “end”: 10,
         “frequency”: 3,
         “conductance”: 0.5e-9,
         “nInputs”: 250,
         “modFile”: “tmGlut”
      }
   }
}
~~~

In the *dSPN* target configuration the *start, end* and *frequency* are specified as vectors. Here the input is 4Hz at 2-3 seconds, and 2Hz at 5-7 seconds. We can also use population units to specify heterogenous external input to the target volume, see `examples/notebooks` on Github.

To create advanced inputs not supported by Snudda the custom spike times can be read from a CSV file (with one spike train per row) by using “generator”: “csv” and “csvFile”:”path/to/your/csvfile”.

A more complex example using additional input generation functionality is given below. The *type* defines what sort of input the synapses form, e.g. *AMPA_NMDA* or *GABA*. The number of inputs to each neuron can either be defined directly using *nInputs* or indirectly by specifying the density of inputs *synapseDensity* along the dendrites. If both parameters are given, the code will use the density but scale it so that the *nInputs* are created. There is also an optional *parameterFile* that can be used to define a set of parameters for the synaptic channel.

The *populationCorrelation* describes how correlated the Poisson input is that is generated by mixing a shared mother process (each spike is included with probability *P* = sqrt(*C*)) and a number of independent child processes (inclusion probability 1 - *P*) to get the resulting input trains (Hjorth et al., 2009).

~~~
{
   “dSPN”: {
      “CorticalSignal” : {
      “generator” : “poisson”,
      “start” : 1.0,
      “end” : 1.5,
      “type: “ : “AMPA_NMDA”,
      “synapseDensity” : “0.05/(1+exp(-(d-30e-6)/5e-6))”,
      “frequency” : 1,
      “populationCorrelation” : 0.0,
      “jitter” : 0.002, “conductance” : 0.5e-9,
      “modFile”: “tmGlut”,
      “parameterFile”:
“$DATA/synapses/striatum/M1RH_Analysis_190925.h5-parameters-MS.json”
   },
   “Thalamic” : {
      “generator” : “poisson”,
      “type: “ : “AMPA_NMDA”,
      “synapseDensity” : “0.05*exp(-d/200e-6)”,
      “frequency” : 1,
      “populationCorrelation” : 0.0,
      “jitter” : 0.01, “conductance” : 0.5e-9,
      “modFile”: “tmGlut”,
“parameterFile”:”$DATA/synapses/striatum/TH_Analysis_191001.h5-parameters-MS.json”
   }
},
~~~

We also include the functionality of virtual neurons, which are neurons that are not simulated, instead their activity is driven by a predefined spike train. This can be used to model, for example, the activation of reconstructed cortical axons in the striatum, which after touch detection will drive the neurons they connect to.

When using synapse density to place excitatory input onto the neurons, larger neurons will receive more input than smaller neurons of the same type. However, size does not necessarily correlate with excitability of the neuron, or steepness of the I-V curve which depends on intrinsic channels. To handle this variation of excitability Snudda allows the user to scale the number of synapses reaching a neuron, a process which in a real network might be regulated by neuronal homeostatic processes.

### What happens under the hood?

For each volume modelled the cell placement is restricted to be inside the mesh specified. The neurons are placed one by one, with coordinates randomly sampled from a uniform distribution. If a neuron position is inside the mesh, and there are no other neurons within a distance *dMin* from it, the position is accepted. To avoid an artificial increase of neuron density at the border, the neuron positions placed outside the mesh are also tracked. These padding positions are not counted towards the total, and are discarded afterwards. Orientation of the neurons in the striatum is completely random, but it is possible to specify other ways to sample the orientation.

For touch detection the space is divided into voxels of 3 μm side length (**Figure 3B**). Synapses are only detected when axon and dendrite are present in the same voxel. The maximal interaction distance is thus decided by the voxel size. To parallelise the touch detection the voxels are grouped into *hypervoxels*, containing 100^3^ voxels each. The mouse dorsal striatum occupies about 26.2 mm^3^ (https://mouse.brain-map.org/), and contains almost 2 million neurons (Rosen and Williams 2001). The first step is to identify which neurons belong to which hypervoxels. For a large portion of the neurons they will be present in more than one hypervoxel. This procedure is done in parallel where the worker nodes of the parallel computer get allocated a subset of the neurons and based on the vertex coordinates of the neurites calculate which hyper voxel the neurons are in. The results are gathered, creating a list of neurons for each hypervoxel. The hypervoxels are then sorted based on the number of neurons inside, and those with most neurons are processed first for better load balance. To perform the touch detection, a line parsing algorithm takes small steps *Δx, Δy, Δz* along all line segments of the dendrites, marking the voxels they intersect. The voxels contained within the soma are also marked. It then repeats the procedure for the axon line segments of the morphologies. Voxels that contain both axons and dendrites are considered to have a putative synapse if the two neuron types are allowed to have a connection between them (**Figure 3A,B**).

The purpose of the touch detection is to find the potential locations where neurons can connect to each other based on the restrictions set by the morphologies. The result of the above touch detection is a set of putative synapses, which is larger than the set of actual synapses. In the pruning step, the set of putative synapses is reduced to match the connectivity statistics from experimental pairwise recordings.

In experiments it is common to report only the binning size and the number of connected neuron pairs, and the total number of pairs. It would be beneficial for circuit modelling if the distance for each pair was also recorded, we could then extract distance dependent connectivity profiles and compare those to what the computer models predict. The pruning is divided into multiple steps, described above. The touch detection for a cubic millimeter can be run in a couple of hours on a desktop, and the whole striatum can be created in a couple of hours on a supercomputer (**Figure 5**). As an example, creating a striatal network with 10,000 neurons (6.4 million synapses and 1468 gap junctions) on a desktop Intel Xeon W-2133 CPU @ 3.60GHz with 6 cores and 64GB RAM took: *init* ∼1 s, *place* 10 s, *detect* 8 min and *prune* 6 min. For 20,000 (50,000) neurons the corresponding times are ∼1 s (∼1 s), 16 s (32 s), 15 min (38 min), 12 min (36 min) to detect 14.3 million (39.9 million) synapses and 3462 (9395) gap junctions.

**FIGURE 5.**
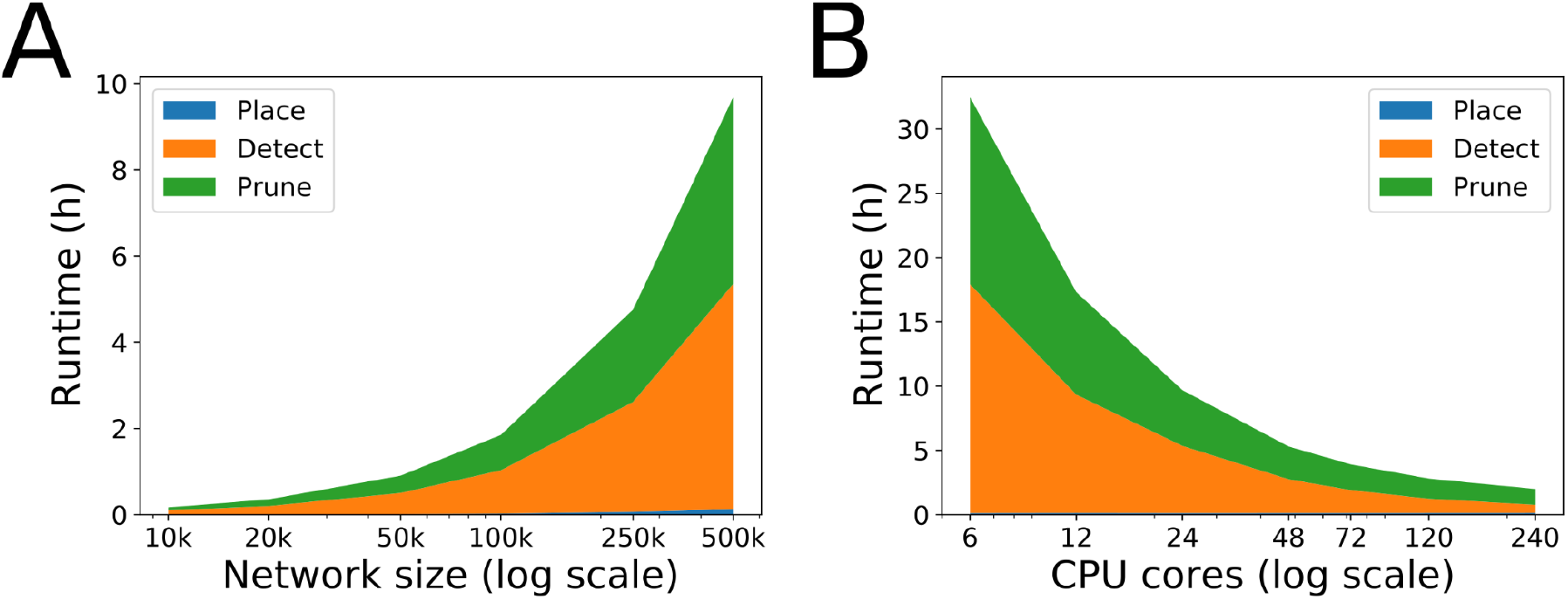
Snudda benchmarking on Tegner cluster at PDC/KTH. Each node has Intel E5-2690v3 Haswell with 2×12 cores and 512 GB RAM. **(A)** Runtime on one node (24 CPU cores) for different network sizes. **(B)** Runtime as a function of the number of CPUs when creating a network with 500,000 neurons, around 469 million synapses and a hundred thousand gap junctions. Place takes very little time compared to the other two phases and is barely visible at the bottom of the two graphs.

In addition to JSON configuration files, the resulting network data is stored in HDF5 files.

### The challenge of limited morphology data

A big challenge of the biologically detailed anatomically constrained simulations of the neural microcircuits is availability of the high quality morphological reconstructions of the main neuron types, in sufficient numbers and variability. Open public morphometric repositories, similar to ModelDB for models present in the world-wide web since 1996 (McDougal et al., 2017), were pioneered in 2006 by G. Ascoli with NeuroMorpho.Org (Akram et al., 2018; http://neuromorpho.org/) and continued by other research centers like Allen Brain Institute (Jones et al., 2009; https://portal.brain-map.org), Janelia Research Campus (Gerfen et al., 2018; Economo et al., 2019; http://mouselight.janelia.org/), eBRAINS Knowledge Graph (https://kg.ebrains.eu/), to name a few, have become increasingly popular among computational neuroscientists.

Single-cell morphological reconstructions vary in quality due to the difference in experimental procedures leading to varying degrees of physical integrity of the neurites, spatial resolution, tissue shrinkage, slicing, etc. Need for consistent quality assurance of morphological reconstructions facilitated development of morphology processing tools for morphometric measurements, data processing and error correction, such as L-measure (Scorcioni et al., 2008; http://cng.gmu.edu:8080/Lm), TREES toolbox (Cuntz et al., 2010; https://www.treestoolbox.org/), btmorph (Torben-Nielsen, 2014; https://bitbucket.org/btorb/btmorph) and NeuroM/NeuroR (Anwar et al., 2009; https://github.com/BlueBrain/NeuroM; https://github.com/BlueBrain/NeuroR). Here we will illustrate typical use cases of manipulating morphological data on the example of a small Python module *treem* (https://github.com/a1eko/treem), developed by the authors in conjunction with Snudda as a complementary instrument to above mentioned packages.

Module Treem provides data structure and command-line tools for accessing and manipulating the digital reconstructions of the neuron morphology in Stockley-Wheal-Cannon format, SWC (Cannon et al., 1998). Access to morphological data from the source code is supported by several Python classes. Common operations with SWC files are possible from the user-written scripts or via the command-line tool `swc`. For the detailed description of the user interface, see API and CLI references in the online documentation (https://treem.readthedocs.io).

A common reconstruction error is so called “z-jump” (Brown et al., 2011) when a part of the neurite gets shifted along the *z*-axis by a few micrometers as shown in **Figure 6A** (top panel). These can result from an accumulated error during the manual reconstruction or as a mistake in automatic procedure. Possible z-jumps can be eliminated in Treem by the `repair` command using one of the four methods, *align, split, tilt* or *join*, as illustrated in **Figure 6A**. Choice of the repair method as well as the assessment of the result should ideally be left to the author of the reconstructed data; if this is not possible the preference is given to the method which better preserves cell symmetry.

**FIGURE 6.**
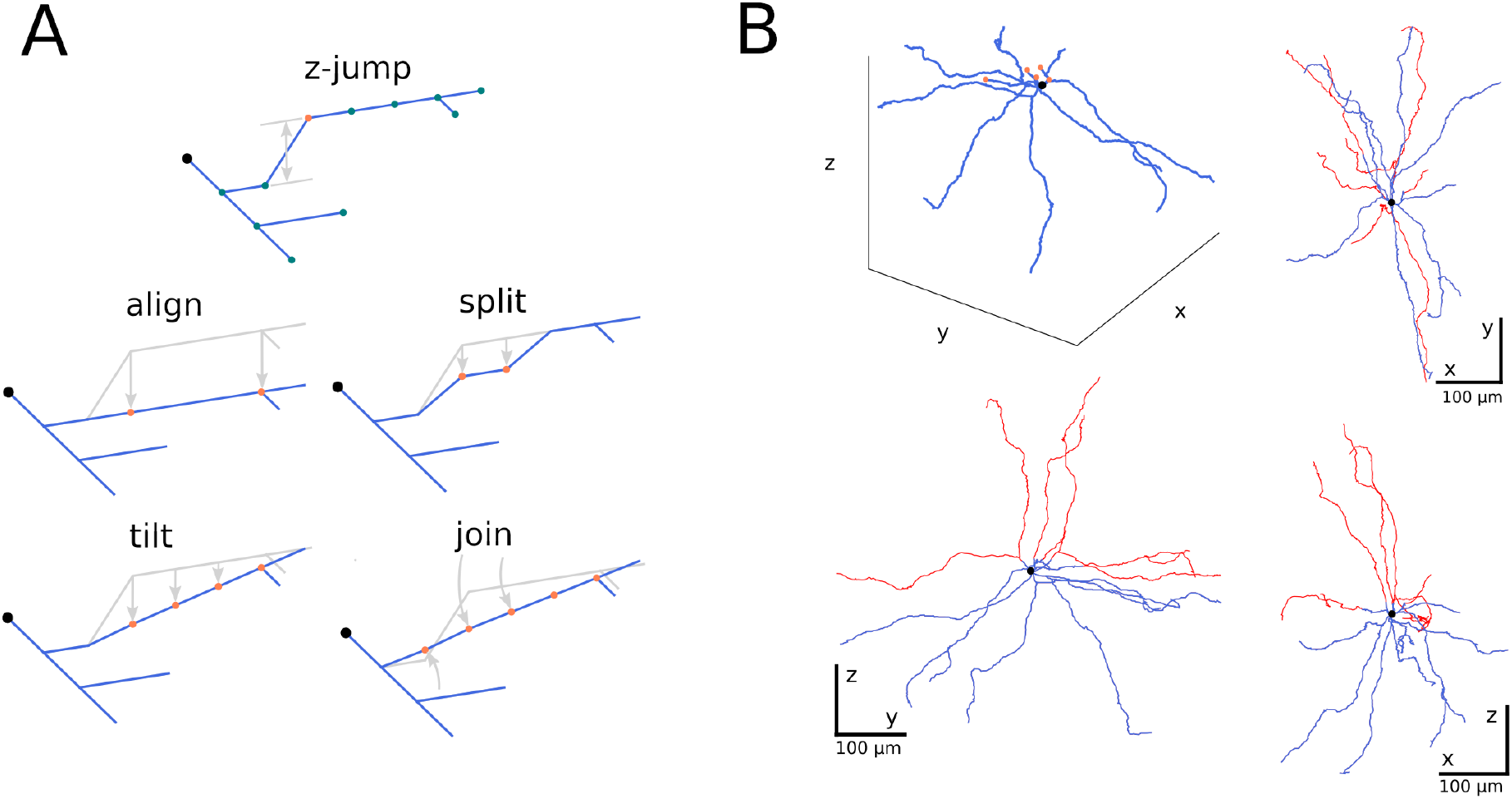
Repairing digital reconstruction of the neuron morphology. **(A)** Correcting “z-jump” reconstruction errors (top panel). Dots illustrate reconstructed points, soma is black, dendrites are blue, the orange dot labels the node at the point of presumed discontinuity. Four correction methods implemented in Treem (Python module *treem*) are shown below (*align, split, tilt* and *join*). **(B)** Repairing the dendrites cut at the slice border. Orange dots in a 3D plot show the cut points of the dendrites. Red lines in 2D projections show “repaired” dendrites, i.e. extended neurites using undamaged reconstructions of the same topological order as the cut branches.

Since neuronal tissue can shrink due to dehydration during histological preparation, correction factors are required before a reconstructed morphology can enter the simulation pipeline. Shrinkage correction involves scaling of the entire reconstruction in (*x, y*)-plane, expansion in *z*-direction, as well as decreasing contraction of selected neurites, e.g. dendrites, by stretching along their principal axes (termed “unravelling” in Markram et al., 2015) or length-preserving spatial filtering (not shown, see online documentation).

Another important omission is that the neurons located close to the slice surface often have their neurites cut and thus missing in the digital reconstruction. Cut neurites can be replaced using the intact branches of the same topological order from the inner part of the slice, assuming spherical or axial symmetry of the neuron morphology as shown in **Figure 6B**. In Treem this is achieved with the `repair` command (see online documentation for the example commands to reproduce **Figure 6B**).

One of the aspects of the large-scale simulations is realistic variability of the model parameters mimicking the natural spread of morpho-electric characteristics in live neurons. To enforce variability in the simulation based on the limited number of reconstructed neurons, we apply random manipulations to the morphological reconstructions. Examples of the length-preserving modifications implemented in Treem are shown in **Figure 7A**. Methods *jitter, twist* and *rotate* do not change the length of the dendritic branches and thus do not affect electrophysiological features of the optimized models but help to recover spatial symmetry of the morphological reconstructions as shown in **Figure 7B**. To distribute excitability of the single-cell models, digital reconstructions can be scaled randomly in 3D, as was done in the large-scale simulation by Hjorth et al. (2020) and illustrated in the online documentation of Treem (https://treem.readthedocs.io).

**FIGURE 7.**
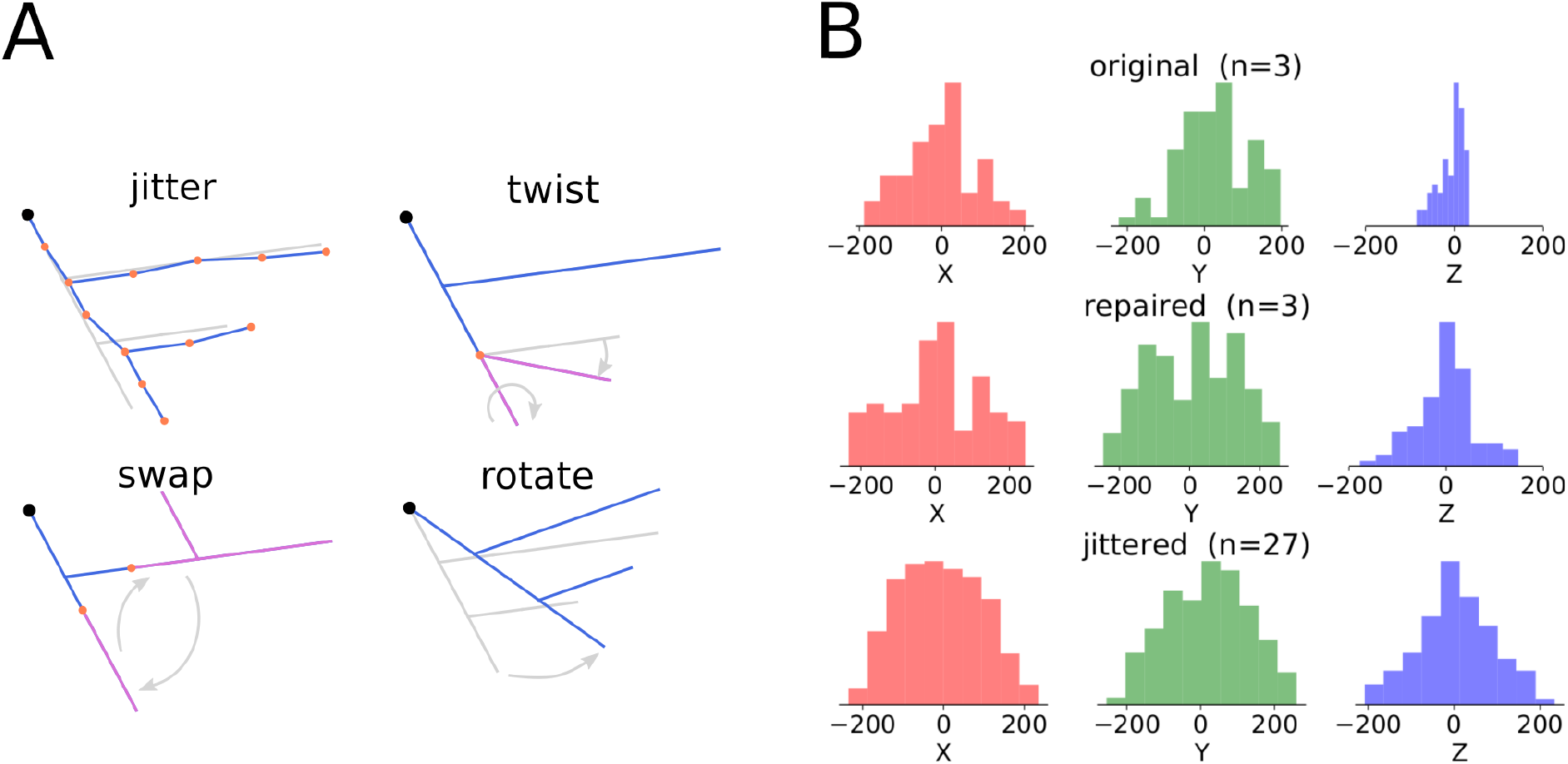
Adding variability to the morphological reconstructions. **(A)** Examples of modification methods used in Treem (modifications preserving the total length are shown). **(B)** Distribution of the coordinates of the dendritic terminations of the fast-spiking interneurons at different stages of morphology processing - original reconstructions (*n*=3), repaired reconstructions (*n*=3) and “jittered”, i.e. randomly manipulated reconstructions, nine variants per each cell (*n*=27). Here, only random twisting of the dendritic branches at the bifurcation points was applied which proved to be sufficient to restore the symmetry.

## Discussion

In this paper we show how to use our open source modelling pipeline to build microcircuit models. An important goal is that the model building process should be transparent and possible to reproduce by other labs, and the model should be extendable when new data accumulate. The pipeline is developed for setting up large-scale simulations of subcortical nuclei, such as striatum. In our current pipeline, based on the software Snudda, neurons can be placed in a defined volume, and then prediction of the location of synapses (as well as gap junctions) can be made using neuron morphologies and the specified pruning rules. Also synaptic data for short-term synaptic plasticity or failure rates can be represented. Finally a simulation using the NEURON simulation environment can be launched. A challenge when using cellular level data from public databases is that sometimes the data for the reconstructed neuronal morphologies only include soma and dendrites, missing the axon entirely. Therefore Snudda supports the prediction of synapses using different approaches. If the detailed morphology is available, the reconstructed axons and dendrites can be used to constrain which neurons are within reach of one another. If the axon is missing, the user can instead specify an axonal density which is then used for the synapse detection. Also our pipeline provides the opportunity to ‘repair’ the dendritic morphologies, and we have illustrated ways to do this using the software Treem. In addition, Treem can provide jittering of morphological parameters to increase the variability in the modelled population of neurons, which is useful to avoid artefacts when there are too few available morphologies for each neuron type. The current version of Snudda does not treat spines separately from the rest of the dendrites, a future improvement would be to allow the user to specify requirements to target spines separately, e.g. if spines are already specified in the reconstruction data.

Models of neocortical microcircuits have been built with similar approaches (see above), however, not all elements of their workflow were available or open-source at the time of Snudda development. We believe that our open source pipeline might become useful when building biophysically detailed microcircuit models of other subcortical brain regions, such as the other basal ganglia nuclei.

When using the current modelling workflow, it is assumed that one has a collection of quantitatively detailed neuron models for each neuron type to be used in the modelled microcircuit. Such neuron models might come from public databases (see above). But most likely several of the neuron types in the selected microcircuit to be modelled might have to be built from scratch. Here the challenges are several as described above. Data on electrophysiological recordings published might be incomplete in such a way that only a few selected traces, as shown in the published manuscripts, exist. Also transcriptional, electrophysiological and morphological data might come from different experiments. Ideally, however, it would be best to have recordings from the neurons that were morphologically reconstructed, such as in patch-seq technique (e.g. Fuzik et al., 2016). Although the electrophysiological properties are well studied, one might lack the knowledge of which ion conductances are expressed in the neurons. Fortunately, such data are starting to emerge, and for example, for striatum transcriptomics data already exist (Muñoz-Manchado et al., 2018; Ho et al., 2018; Gokce et al., 2016; Saunders et al., 2018). If one has a good hypothesis of which channels are expressed, characterisation as well as models are starting to be collected at resources such as the Channelpedia (Ranjan et al., 2011; http://channelpedia.net) and Ion Channel Genealogy (Podlaski et al., 2017; https://icg.neurotheory.ox.ac.uk/). Although still not trivial, if one has both the morphology and electrophysiological data of a particular neuron type, workflows have already been developed for optimizing neuron models (Van Geit et al., 2016; Migliore et al., 2018; Masoli et al., 2020).

A natural future goal would, however, be to link microcircuit models built in different labs, e.g. a cortical microcircuit connected to a striatal microcircuit. Then interoperability between models as well as model specification, such as SONATA, would be crucial. We have on our road map for Snudda to support the SONATA standard (Dai et al., 2020) and work has already started on it to become interoperable with the EBRAINS infrastructure (https://ebrains.eu/).

## Acknowledgements

The simulations were performed on resources provided by the Swedish National Infrastructure for Computing at PDC (Center for Parallel Computing). We acknowledge the use of Fenix Research Infrastructure resources, which are partially funded from the European Union’s Horizon 2020 research and innovation programme through the ICEI project under the grant agreement No. 800858. The authors wish to thank Sten Grillner, Johanna Frost-Nylén, Robert Lindroos and Ilaria Carannante for helpful discussions. We also thank Robin de Schepper, Kadri Pajo, and Wilhelm Thunberg for help with software compatibility.

## Declarations

### Availability of data and material

Data used for single-cell neuron models as well as synaptic connectivity within the example network of the striatal microcircuit is available on GitHub (in the `snudda/data` and `snudda/examples` folders at https://github.com/hjorthmedh/Snudda).

### Code availability

(see Information Sharing Statement)

### Information Sharing Statement

The presented software Snudda (version XXX; RRID:SCR_021210) and Treem (version YYY, DOI:YYY) are available on GitHub and PyPI:

- Snudda - https://github.com/Hjorthmedh/Snudda, https://pypi.org/project/snudda/
- treem - https://github.com/a1eko/Treem, https://pypi.org/project/Treem/

